# Balancing Inhibition and Sparsity for Stable, Accurate Cerebellar Learning

**DOI:** 10.64898/2026.04.09.717611

**Authors:** Liao Yu, Zhuoqin Yang, Yeyao Bao, Yunliang Zang

**Affiliations:** School of Mathematical Sciences, Beihang University, Beijing, 100191, China; Academy of Medical Engineering and Translational Medicine, Medical Faculty, Tianjin University, Tianjin, 300072, China

## Abstract

The cerebellum’s structured circuitry supports learning across motor and cognitive domains, yet the coding strategies in granule cells that enable this versatility remain unclear. Using a theoretical cerebellar model, we examined how feedforward inhibition (FFI) and feedback inhibition (FBI) shape granule cell activation patterns to optimize learning in two tasks: complex trace learning and pattern identification. For trace learning, performance depends on spatiotemporally ordered granule cell activity shaped by FBI, with temporal sparsity emerging as the key determinant of accuracy. For pattern identification, both pathways support high accuracy, with only slight sensitivity to pathway choice. In both tasks, spatial sparsity becomes critical in incremental learning to prevent memory interference, a role reinforced by advanced synaptic plasticity strategies. These findings identify sparse granule cell activation as a unifying principle for cerebellar learning and reveal task-dependent roles of inhibitory pathways, providing a mechanistic framework for understanding stability–plasticity trade-offs in cerebellar learning.

## INTRODUCTION

The cerebellum’s compact circuitry hides remarkable computational versatility— capable of guiding both precise motor actions and abstract cognitive functions. Traditionally recognized for motor coordination and sensorimotor learning, the cerebellum is now implicated in diverse cognitive functions, including pattern separation, mental arithmetic, and decision-making ^1-8^. This breadth of involvement raises a fundamental question: how can a relatively simple and highly structured circuit supports such universality at the computational level?

Cerebellar circuitry is organized such that mossy fibers convey sensory or contextual inputs to granule cells, which recode the information and relay it to Purkinje neurons for output ^1,9-12^. Granule cells—the most numerous neurons in the brain—are central to cerebellar computation, providing a high-dimensional basis for transforming inputs into precise motor trajectories or abstract classifications. A critical feature of granule cell processing is the presence of two distinct inhibitory pathways: feedforward inhibition (FFI), in which mossy fiber input excites Golgi cells that inhibit granule cells, and feedback inhibition (FBI), in which granule cell activity excites Golgi cells that inhibit granule cells in return ^13-19^. These pathways differ in timing and spatial targeting, potentially influencing both the diversity and sparsity of granule cell firing patterns. Yet, their relative contributions to learning remain unresolved: they might be redundant, complementary, or selectively recruited depending on task demands.

Sparse coding in cerebellar granule cells, as envisioned in the Marr–Albus theory, is thought to boost learning efficiency and minimize interference by maximizing pattern separation ^9,10,14,20-22^. Direct experimental confirmation remains elusive due to technical constraints. Calcium imaging in zebrafish and mice indicates dense granule cell activation ^23-25^, yet the slow kinetics of calcium indicators obscure precise temporal patterns and underestimate silent cell fractions ^26^. Evidence from other studies ^27^ suggests that true sparsity may be concealed, underscoring the need to clarify how granule cell dynamics support efficient learning.

Moreover, the optimal activation patterns—and their dependence on inhibitory pathway choice—remain unknown. Previous theoretical studies have examined cerebellar learning in motor tasks ^28,29^ or pattern separation ^9,21,30,31^, but rarely within a unified framework that allows direct comparison. Most models also focus on acquiring a single task, neglecting the biological requirement for incremental learning ^32,33^—acquiring new knowledge without erasing old memories. The mechanistic link between inhibitory balance, coding sparsity, and the stability–plasticity trade-off has not been systematically explored.

Here, we address these gaps using a theoretical cerebellar circuit model with tunable inhibitory balance, enabling precise control over granule cell activation levels and spatiotemporal firing patterns. We examine two distinct tasks—complex trace learning and pattern identification—under both single-task and incremental learning conditions. This dual-task approach allows us to identify shared principles, such as granule cell activation sparsity, and task-specific differences in pathway sensitivity. Specifically, we ask: How do FFI and FBI pathways differentially shape granule cell activity and learning performance? What forms of sparsity—temporal *vs*. spatial—optimize learning, and how do these differ between single-task and incremental learning? Are these principles general across motor and non-motor tasks, or does pathway sensitivity depend on task demands?

We find that granule cell sparse activation is a unifying coding strategy across motor and non-motor cerebellar tasks and reveal task-dependent roles of inhibitory pathways. Complex trace learning depends on spatiotemporally ordered activity shaped by the FBI pathway, whereas pattern identification remains robust under both pathways. Spatial sparsity emerges as critical in incremental learning to prevent memory interference, linking inhibitory balance and coding density to stability–plasticity trade-offs.

## RESULTS

We constructed a cerebellar circuit model (Figure 1) that captures the essential feedforward flow from mossy fibers to granule cells to Purkinje neurons. The balance between FFI and FBI was controlled by α (α = 0: pure FBI; α = 1: pure FFI). This architecture enables direct testing of how inhibitory balance and coding sparsity shape learning performance across tasks with different temporal and representational demands.

**Figure 1.**
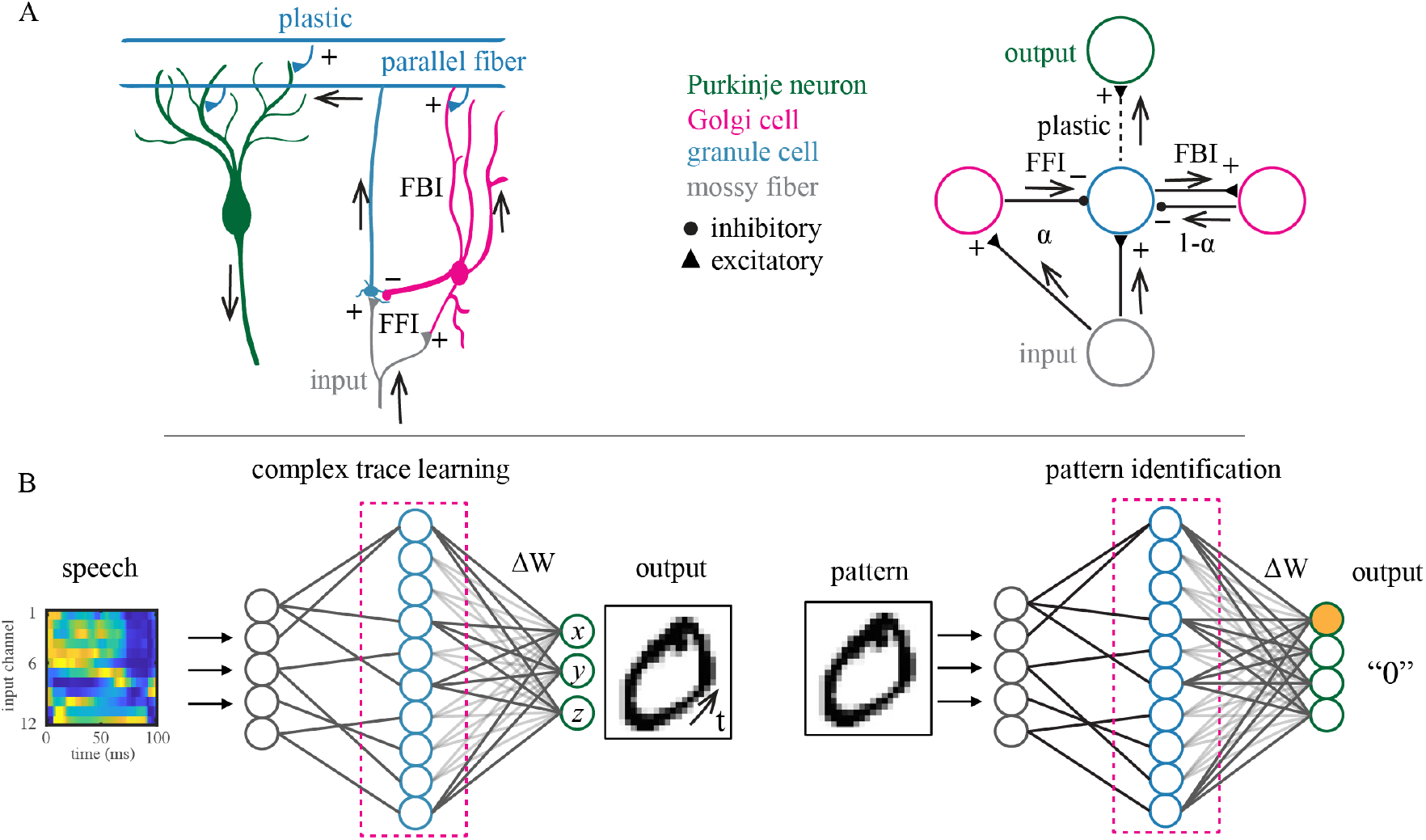
Schematic of Cerebellar Model and Cerebellum-associated Tasks. (A) Simplified cerebellar architecture. Mossy fibers project to granule cells, which recode input and relay it to Purkinje neurons. The mossy fiber → Golgi cell → granule cell pathway forms FFI, and the granule cell → Golgi cell → granule cell pathway forms FBI. Parameter α controls their relative influence. (B) Two tasks: complex digit trace learning (left), where the model predicts temporal *x*–*y*–*z* coordinates from a spoken digit spectrogram, and pattern identification (right), where the model predicts input identity from an MNIST image pattern. The dashed box indicates regulation of granule cell activity by FFI and FBI.

We trained the model on two tasks: complex trace learning from spoken inputs (TI-46 corpus ^34^) to mimic sensorimotor transformation, and image classification (MNIST ^35^) to mimic pattern identification, under both single-task and incremental learning conditions. All synapses were fixed except for plastic granule cell → Purkinje neuron connections, which were updated using gradient descent. We also tested advanced synaptic update methods—elastic weight consolidation ^36^ and synaptic intelligence ^37^. Detailed model architecture, parameters, and training procedures are described in the Method Details section.

With this framework established, we next examine how inhibitory pathway balance and granule cell activation patterns shape learning performance in each task.

### Complex Trace Learning Relies on FBI Pathway

Sensorimotor learning requires transforming sensory inputs into continuous motor trajectories in spatial coordinates, a process that depends critically on how information is dynamically represented between input and output layers. Both FFI and FBI can influence the diversity and sparsity of this representation. To assess their relative roles in shaping granule cell activity and learning performance, we challenged the cerebellar model with predicting the complex spatiotemporal trace of a single digit.

Simulation results show that learning performance is strongly determined by the source of inhibition. Both granule cell activation level and inhibitory pathway balance, quantified by α, substantially affect prediction performance (Figure 2A). Performance improves as α decreases, reaching its lowest error when granule cell activation is shaped exclusively by FBI (α = 0). In contrast, pure FFI mediation (α = 1) fails to predict any trace accurately, even for the simplest line-like digit “1” (Figure 2B). Across all digits tested, learning error increases monotonically with α (Figure 2C).

**Figure 2.**
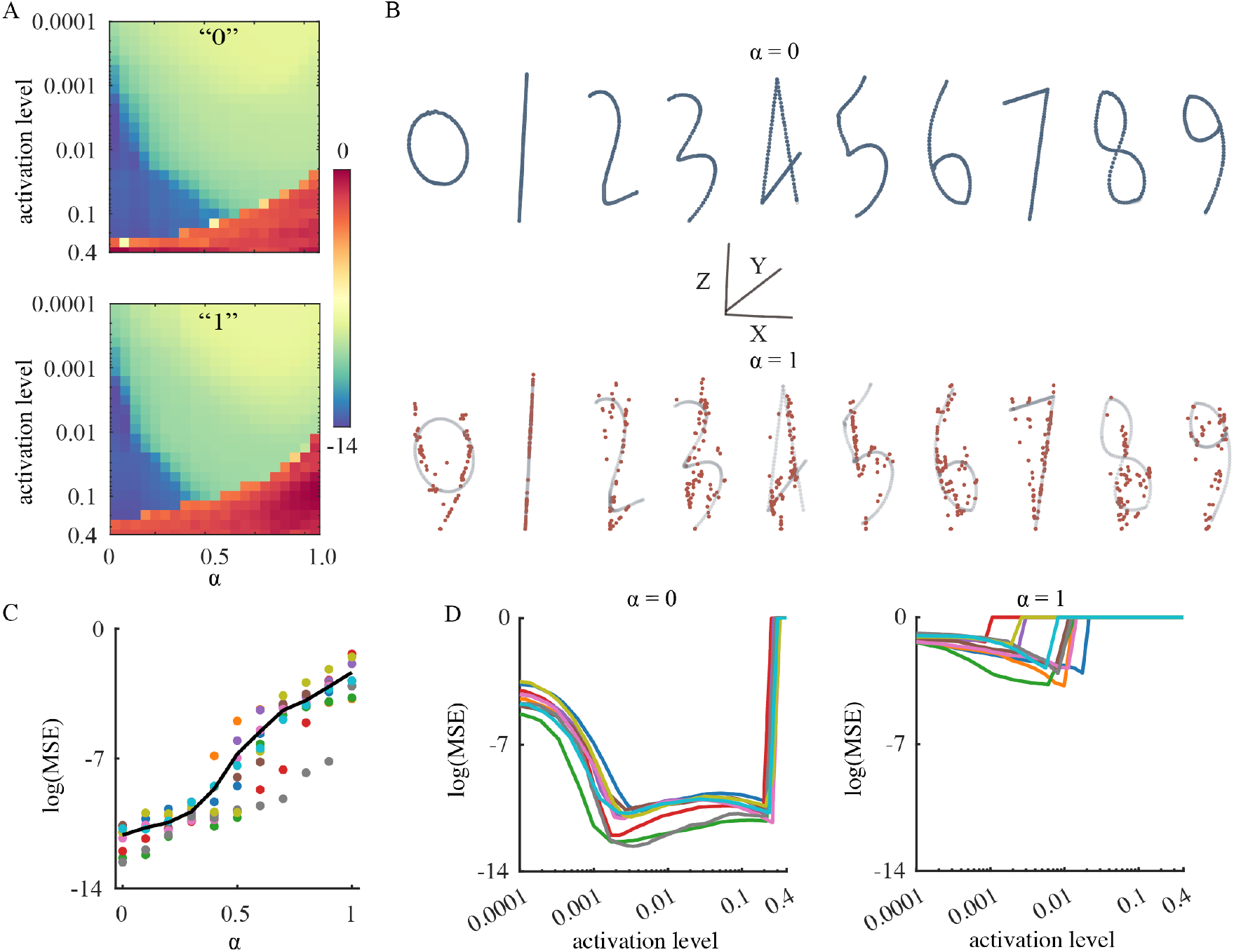
Single-digit Complex Trace Learning. (A) Heat maps of mean squared logarithmic error, log(MSE), for digits “0” (top) and “1” (bottom) across inhibition balance (α) and granule cell activation levels. (B) Predicted traces with α = 0 (FBI) and α = 1 (FFI). (C) Learning error versus α. (D) Learning error versus activation level for α = 0 and α = 1. In C-D, colors indicate individual digits.

For the FBI-mediated model, optimal performance occurs within a relatively sparse granule cell activation range (in the range of 0.001 and 0.2), whereas both lower and higher activation levels significantly increase error. In the FFI-mediated model, prediction errors remain consistently high across all activation levels. These findings demonstrate that FFI and FBI regulate granule cells in distinct ways, with FBI providing patterns for accurate sensorimotor learning, while FFI fails to support trace prediction.

### Granule Cell Activation Patterns Differ by Inhibition Pathway

To delineate the distinct modulatory effects of FFI and FBI on learning performance, we analyzed the spatiotemporal patterns of granule cell activation. Under identical input conditions, these patterns differ markedly depending on which inhibitory pathway mediates the response (Figure 3).

**Figure 3.**
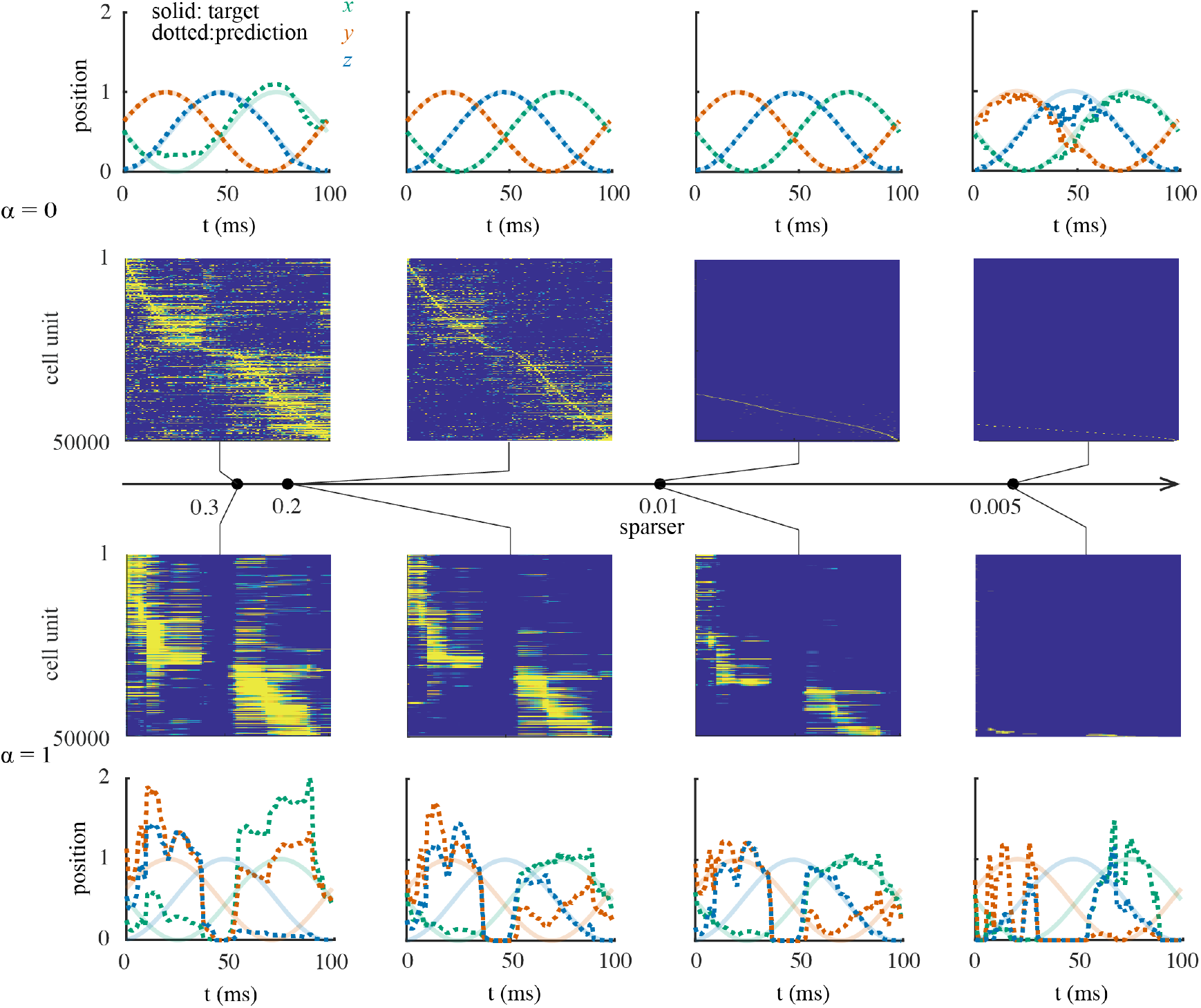
Inhibition Pathways Shape Granule Cell Activation and Cerebellar Learning. Granule cell activation patterns (second and third rows) and corresponding learning outcomes (first and fourth rows) under pure FBI (top) and pure FFI (bottom) mediation for digit “0” learning. Inhibition strength increases from left to right, resulting in progressively sparser activation of granule cells.

Under pure FBI mediation, granule cell responses form distinct, time-ordered sequences across the population at all activation levels, consistent with previous observations in cerebellar granule cells of mice and in the electrosensory lobe of electric fish ^25,38^. As inhibition strengthens and activation levels decrease, both spatial (population-level) and temporal (lifetime activity of individual neurons) activation become progressively sparser. Learning accuracy improves with moderate sparsification but declines when activation becomes excessively sparse.

In contrast, FFI produces temporally dense activity—prolonged periods of activation followed by extended pauses—and spatially synchronized firing. As with FBI mediation, stronger inhibition reduces both spatial and temporal activation. However, the temporally dense and synchronized pattern under FFI limits the diversity of spatiotemporal information available for learning.

Accurate transformation of continuous sensory inputs into complex, time-varying trajectory outputs requires granule cells to provide a rich spatiotemporal basis set. The silent periods following clustered activation during FFI mediation only partially account for its failure in trace prediction, as seen in segments aligned with these pauses. Even during intervals of active granule cell firing, spatial trajectories are poorly learned, indicating that factors beyond silent periods contribute to prediction failure.

To identify the specific features of granule cell activation that optimize complex trace learning, we next designed controlled firing patterns to systematically dissect their contributions.

### Temporal Sparsity as the Key Determinant of Single-Trace Learning

Motivated by theoretical considerations and experimental observations ^25,38-41^, we designed four groups of artificial firing patterns to identify the granule cell activation dynamics that optimize complex trace learning (Figure 4A). The first group (“sync 1– 4”) consists of temporally dense and spatially synchronized firing patterns, with “sync 3” resembling activity mediated by the FFI pathway. The second group (“bio 1–4”) mimics FBI-mediated activity with progressively sparser spatiotemporal activation, reminiscent of experimentally observed granule cell responses ^25,38^. The third and fourth groups (“st 1–7”) comprise seven spatiotemporal patterns specifically designed to isolate the effects of temporal and spatial sparsity. In “st 1–3,” each neuron fires only once during the learning window, while the number of transiently active granule cells gradually decreases. In “st 5–7,” exactly one granule cell is active at any given time point, but each neuron’s activation frequency progressively decreases. The “st 4” pattern combines both conditions—each neuron fires only once, and exactly one neuron is active at any given time.

**Figure 4.**
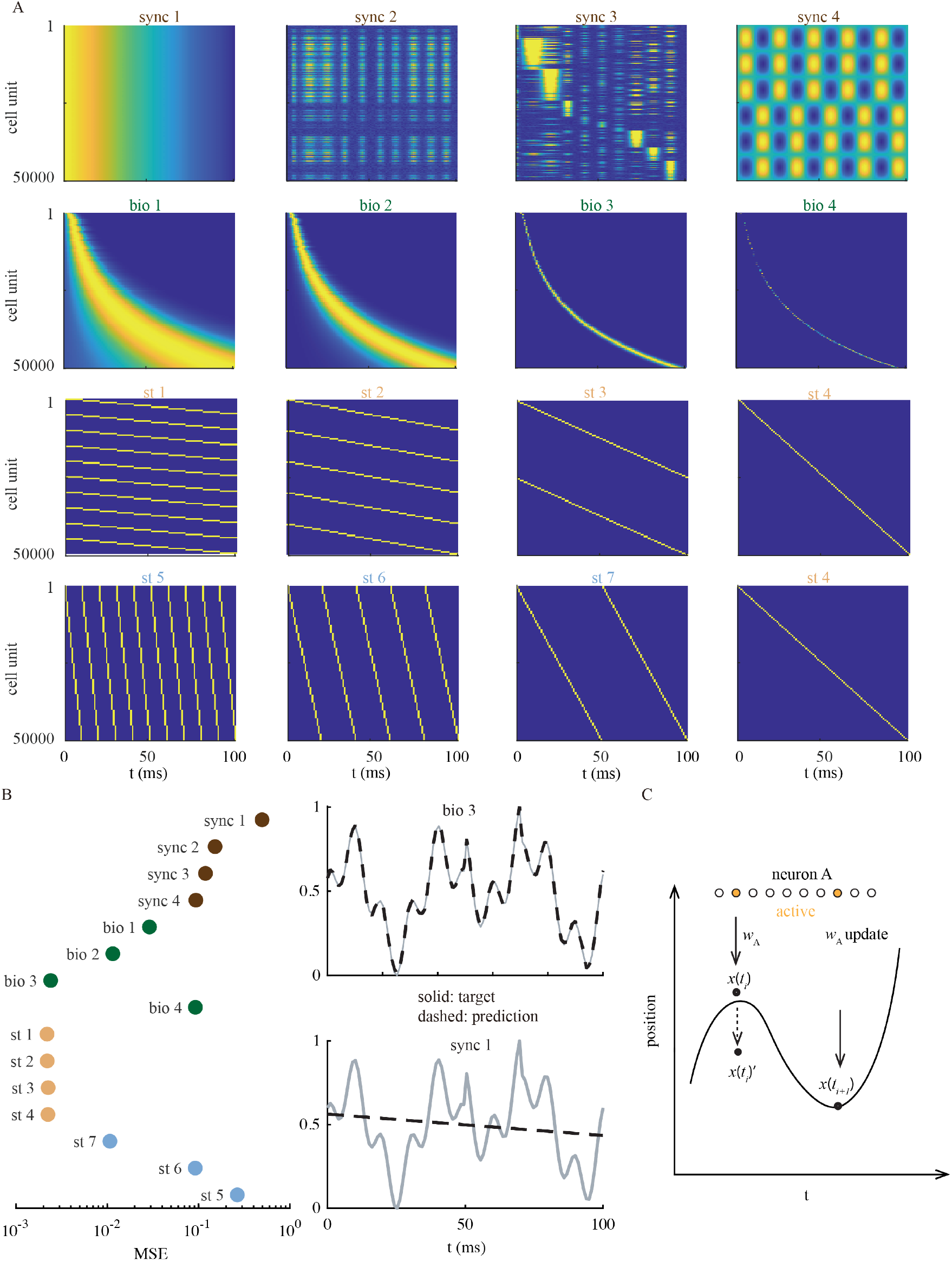
Learning Performance with Synthesized Activation Patterns. (A) Four groups of activation patterns. *Synchronized and temporally dense*: “sync 1”, globally synchronous elevated-then-decayed firing; “sync 2”, globally synchronous alternating firing; “sync 3”, locally synchronous alternating firing; “sync 4”, anti-phased oscillatory firing. *Biologically realistic*: “bio 1–4”, FBI-like activity with progressively sparser spatiotemporal activation. *Temporally sparse*: “st 1–3”, each neuron fires once, with decreasing population activation levels. *Spatially sparse*: “st 5– 7”, constant transient population activation but decreasing firing frequency per neuron; “st 4” combines extreme temporal and spatial sparsity. (B) Learning errors for each pattern (left) and example learning traces (right). (C) Schematic of memory drift caused by neuronal repeated activation.

The cerebellar model fails to achieve satisfactory performance when granule cells fire in clustered, synchronized patterns (“sync 1–4,” Figure 4B). For biologically realistic “bio 1–4” patterns, learning error decreases as activation becomes sparser, reaching a minimum at “bio 3,” beyond which excessive sparsity increases error—likely due to insufficient active neurons to represent inputs. Because both temporal and spatial sparsity increases simultaneously from “bio 1” to “bio 3,” the relative contribution of each factor remains unclear.

This ambiguity was resolved by comparing “st 1–7.” In “st 1–4,” neurons are temporally sparse but population activation levels vary, producing no significant difference in learning performance—indicating that spatial sparsity has minimal impact. In contrast, in “st 5–7,” population activation remains constant while temporal sparsity increases, consistently improving learning accuracy. These results demonstrate that temporal sparsity is the key determinant of accurate single-trace learning, whereas spatial sparsity plays a negligible role.

During trace learning, trajectories evolve continuously over time (Figure 4C). Suppose neuron A is activated to encode position *x*(*t*_*i*_) at time *t*_*i*_. If neuron A is reactivated at *t*_*i*+1_, it must encode a new position *x*(*t*_*i*+1_). When *x*(*t*_*i*_) ≠ *x*(*t*_*i*+1_), the previously learned synaptic weight *w*_*A*_ must be updated to accommodate the new position, introducing memory interference within a single learning episode and causing the original representation of *x*(*t*_*i*_) to drift toward *x*(*t*_*i*_)′.

### Spatial Sparsity Becomes Essential in Incremental Trace Learning

A previous theoretical study suggested that complex trace learning benefits from relatively dense granule cell activity ^29^. In our single-digit trace learning simulations (Figure 2), however, the model achieved optimal performance at extremely sparse activation levels (∼0.001), while errors remained consistently low below 0.2. This apparent discrepancy raises a critical question: during sequential learning of multiple digit traces, what level of granule cell activation minimizes memory interference while preserving the capacity for incremental learning?

Efficient incremental learning requires both stability—minimal overlap among granule cells representing different digits—and plasticity—sufficient representational richness for accurate input–output transformation. Under FBI mediation, optimal learning occurs within the sparse activation range across all task loads (Figure 5A). As the number of learned digits increases, the optimal activation level becomes progressively sparser, while MSE rises with task load.

**Figure 5.**
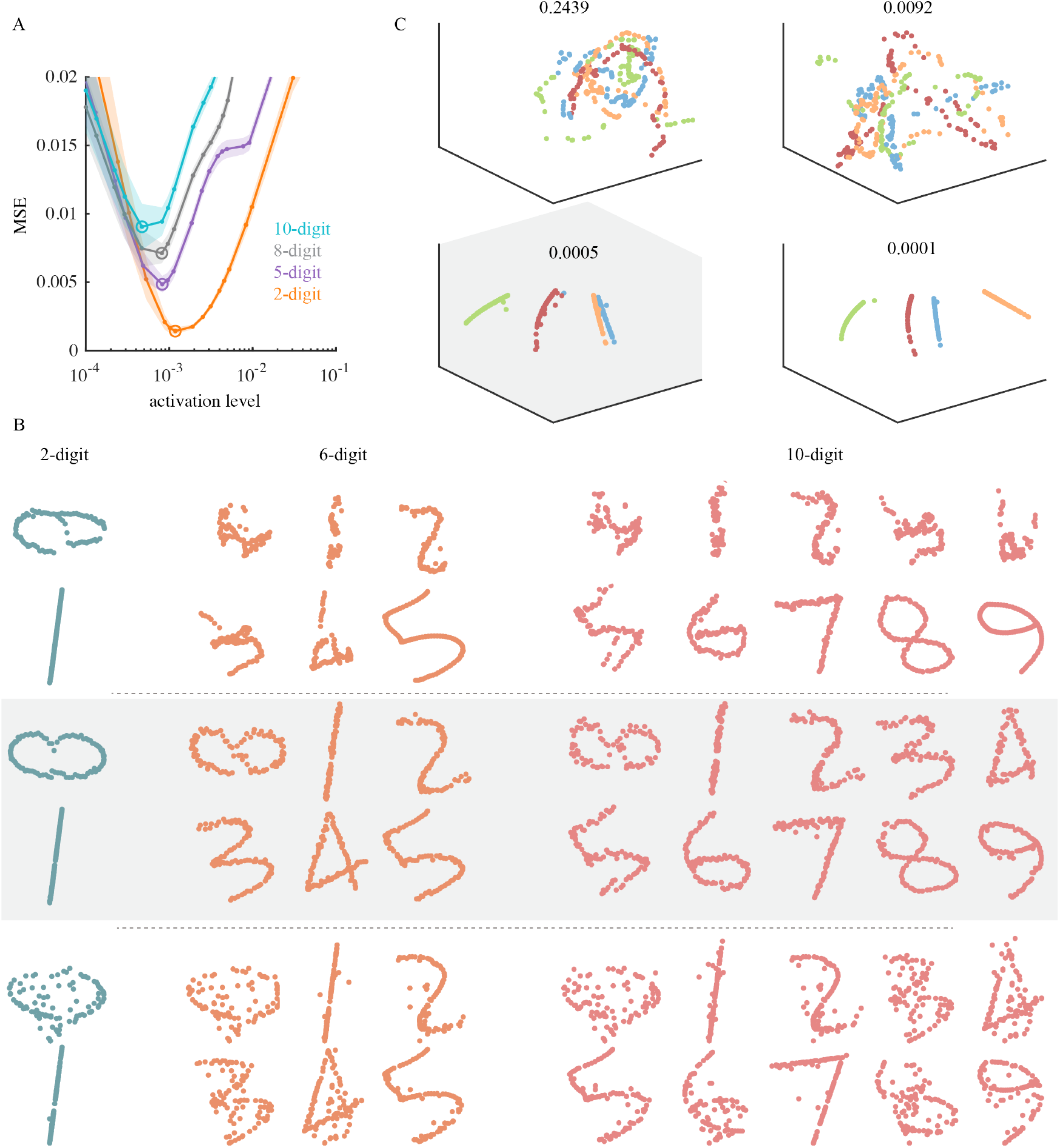
Incremental Trace Learning. (A) Learning error as a function of granule cell activation level. Colors indicate the number of traces learned. (B) Example predicted trajectories after learning 2, 6, and 10 digits, shown for progressively sparser granule cell activation levels (top to bottom). (C) Three-dimensional principal component analysis (PCA) plots of granule cell population activity for four target traces (“1,” “3,” “5,” and “7”) at different granule cell activation levels.

Dense activation provides rich spatiotemporal information and high final-stage accuracy but severely disrupts previously learned traces (Figure 5B), as evidenced by overlapping and poorly defined PCA clusters (Figure 5C). Sparse activation prevents interference by generating distinct neuronal trajectories while retaining sufficient information for new learning (Figure 5B); however, excessive sparsity limits representational capacity. Thus, the optimal activation level balances stability for preserving prior memories with plasticity for acquiring new ones.

While single-trace learning relies primarily on temporal sparsity within individual granule cells (Figure 4), incremental learning additionally requires spatial sparsity to prevent interference. As shown in Figure 6A, dense activation corrupts previously learned traces even when new predictions remain accurate, whereas sparse activation mitigates interference but reduces representational richness. The minimal error occurs at approximately 0.05% activation, with errors increasing at lower activation levels.

**Figure 6.**
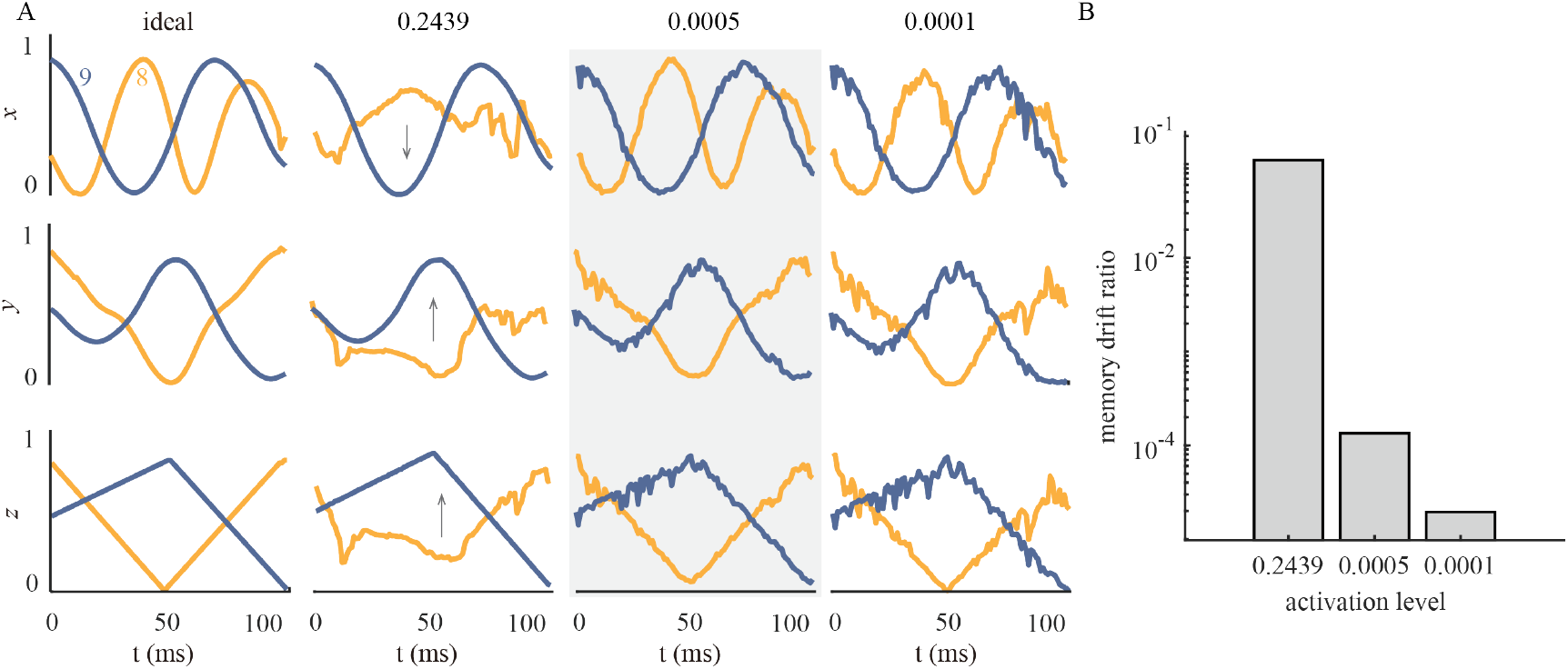
Memory Interference during Incremental Trace Learning. (A) Orange traces represent the first learned digit trace “8,” and blue traces represent the subsequently learned digit trace “9.” The first column shows trajectories from single-digit learning. The second to fourth columns show trajectories for incremental learning with varying granule cell activation levels. Rows correspond to *x, y*, and *z* trajectory components, respectively. (B) Reduced memory drift ratio with sparser activation of granule cells.

To quantify interference, we identified granule cells that were co-activated during the entire learning window for two consecutive digits. For each shared neuron, activity *a*_(*i,j*)_ (*i* = 1, …, *N*) is multiplied by its synaptic weights *w*_(*i,j*)_ during learning of the *j*_*th*_ and (*j* + 1)_*th*_ digits. The *memory drift ratio* for digit *j* after learning digit *j* + 1 is defined as:

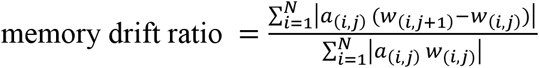

Lower granule cell activation levels produce smaller memory drift ratios, reflecting both enhanced temporal sparsity and reduced overlap between granule cells representing digits *j* and *j* + 1 (Figure 6B).

### Advanced Synaptic Strategies Reinforce the Role of Spatial Sparsity

The importance of spatial sparsity is further supported by advanced synaptic update mechanisms. Elastic weight consolidation ^36^ and synaptic intelligence ^37^ minimize interference by constraining synaptic changes in commonly activated neurons that are essential for retaining prior memories, without affecting temporal sparsity. Both strategies enhance prediction accuracy and reduce memory drift ratios across a wide range of granule cell activation levels (Figures S1, S2). Together, these findings provide convergent evidence that spatial sparsity is indispensable for preventing memory interference and maintaining accurate performance during incremental learning.

### Pattern Identification Depends Weakly on Pathway Choice but Critically on Sparse Coding

In the previous sections, we examined single and incremental complex trace learning. We next asked whether the effects of inhibitory pathway and coding level observed in those tasks generalize to a non-motor pattern identification task. Unlike trace learning, which requires generating spatiotemporal trajectories, pattern identification involves associating static image inputs with discrete outputs ^1^. To test this, we used the widely adopted MNIST dataset to evaluate how feedforward (FFI) and feedback (FBI) inhibitory pathways support pattern identification.

We first examined interleaved learning. In contrast to complex digit trace learning— where FFI mediation fails—both FFI and FBI pathways support high identification accuracy (Figure 7A). Across varying numbers of learned patterns, accuracy increases slightly with α, indicating a modest trend toward improved performance under FFI mediation. Although accuracy declines as the number of learned patterns increased, both pathways maintain high performance across a broad range of granule cell activation levels (Figure 7B). These results suggest that, for relatively simple identification tasks learned in an interleaved manner, the cerebellar model is largely insensitive to inhibition pathway choice and granule cell activation level.

**Figure 7.**
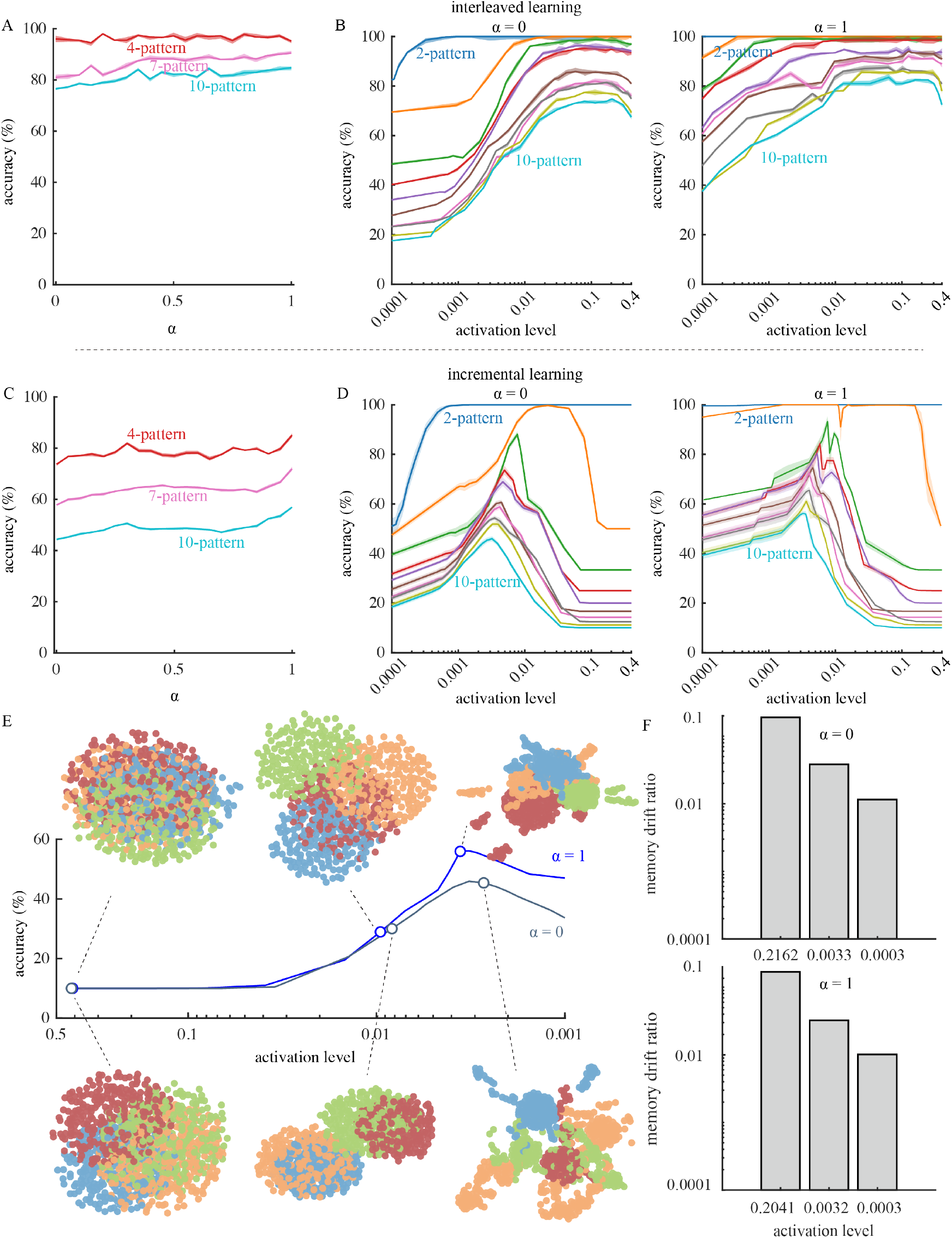
Incremental Pattern Identification Requires Granule Cell Sparse Activation. (A) Maximum identification accuracy as a function of α. (B) Identification accuracy versus granule cell activation level (left: pure FBI mediation; right: pure FFI mediation) for interleaved learning. (C–D) Same as A and B, respectively, but for incremental learning. In A–D, colors indicate the number of learned patterns. (E) PCA scatter plots of granule cell population activity at varying activation levels under pure FFI (top) and pure FBI (bottom) mediation for patterns “1,” “3,” “5,” and “7” during incremental learning. (F) Reduced memory drift ratio with sparser activation of granule cells under the mediation of FBI (top) and FFI (bottom).

In incremental learning, identification accuracy drops markedly compared to interleaved learning. Regardless of the number of learned patterns, accuracy again increases slightly with α, showing a consistent trend toward higher performance under FFI mediation (Figure 7C). As the number of learned patterns increases, the optimal activation range narrows and shifts toward sparser levels (Figure 7D). PCA reveals that sparser activation transforms overlapping pattern clusters into clearly separated groups (Figure 7E). FFI-mediated activity patterns tend to produce more distinct separations than FBI-mediated ones, consistent with the modest advantage in accuracy observed under FFI mediation.

The role of spatial sparsity in reducing memory interference during incremental learning is further confirmed in pattern identification. Regardless of the inhibitory pathway, the memory drift ratio decreases with sparser granule cell activation (Figure 7F). Moreover, both elastic weight consolidation and synaptic intelligence enhance pattern identification by restricting synaptic modifications in neurons commonly activated across patterns and critical for their representation (Figures S3, S4). Together, these results reinforce the importance of sparse coding—particularly spatial sparsity—in maintaining memory stability and accurate performance in static pattern identification tasks.

## DISCUSSION

Using a theoretical cerebellar circuit model, we addressed fundamental questions about the roles of inhibitory pathways, activation patterns, and coding sparsity in granule cells during cerebellar learning. Our results identify sparse granule cell activation as a unifying coding principle and reveal how inhibitory balance adapts to task demands, linking circuit mechanisms to the stability–plasticity trade-off.

Degeneracy and redundancy are defining features of neural circuits, enabling flexibility and resilience in computation ^42,43^. In the highly structured cerebellum, granule cell activity is regulated by two sparsification pathways—FFI and FBI. Although their precise roles remain unresolved experimentally, our findings suggest that these dual routes are central to the cerebellum’s versatility across motor, cognitive, and potentially emotional domains ^1-5^. FFI and FBI are not merely redundant; they can be selectively recruited to match task-specific requirements or reflect regional specializations ^4,26^. This interpretation aligns with evidence that cerebellar Golgi cells are engaged in stimulus- and pathway-specific manners ^27,44,45^. In some contexts, optimal performance may depend on their coordinated action, paralleling cerebellum-like systems such as the fruit fly olfactory circuit, where spike-frequency adaptation and lateral inhibition synergistically sharpen odor discrimination ^46^. Such convergence points to a broader principle: multiple inhibitory pathways can be orchestrated to meet diverse computational needs in hub regions like the cerebellum.

Our results also contribute to the long-standing debate over cerebellar coding strategies, first articulated in the Marr–Albus theory ^14,22^. This debate has been complicated by conflicting calcium imaging results ^23-25,27^ and methodological constraints that have prevented definitive experimental resolution. Here, we show that both spatial and temporal sparsity are essential for incremental learning, whereas dense coding remains effective for specific tasks such as single-trace learning and interleaved pattern identification ^21,29,47^. Sparsity also offers substantial metabolic advantages ^48^: activating most granule cells—which constitute over half of all brain neurons—even for simple behaviors would impose unsustainable energetic costs. Mechanistically, sparsity limits memory interference by reducing representational overlap, whereas multiplexed coding ^49-51^, although potentially increasing computational capacity, risks compromising memory stability. Nevertheless, the functional value of sparsity must be interpreted cautiously, as its benefits are likely contingent on physiologically relevant timescales ^1,26^.

In sensorimotor learning, a well-distributed temporal basis set is essential for representing dynamic positions over time. Granule cells in the cerebellum-like electrosensory lobe of electric fish exemplify this principle, with individual cells responding at distinct moments evenly spaced across behaviorally relevant intervals ^38^. Similarly, in mice trained for forelimb movements, many granule cells display heterogeneous onset and peak Ca^2+^ activity, collectively spanning the entire task duration despite challenges in estimating activation periods ^25^. Remarkably, our simulations show that FBI-shaped granule cell activity mirrors these experimental observations. Our contributions are twofold: (1) we propose a biologically plausible mechanism that accounts for these activation patterns, and (2) we demonstrate that such dynamics optimize complex trace learning.

Most established cerebellar theories focus on motor learning and pattern separation. Early motor studies emphasized eyelid conditioning ^39^ and the vestibulo-ocular reflex ^52,53^, while more recent work has examined complex trace learning ^28,29,47^. Pattern separation theories highlight sparse connectivity and structural properties as critical for discrimination ^1,9,10,30^. However, biologically realistic incremental learning has not been addressed in previous theoretical models. Prior studies have explored consolidation strategies ranging from fast-learning plasticity to slower downstream plasticity ^54^. Here, we propose that moderate sparsity achieves an optimal balance between stability and plasticity. Our unified framework extends incremental learning to both sensorimotor-relevant complex trace learning and cognition-related pattern separation, enabling analysis of task-dependent coding strategies and inhibitory pathway recruitment within a single model.

As evidence grows for cerebellar involvement in diverse tasks ^1-6^, its circuit-level implementation of cognitive functions remains unclear. Our framework provides a foundation for extending cerebellar modeling to additional cognitive domains and clarifying task-dependent granule cell representations. We also offer testable predictions: depressing Golgi cell activity or blocking its inhibition onto granule cells should increase coding density ^27^ and impair learning of complex movements. While this study employed gradient descent and other advanced synaptic strategies that theoretically approximate the upper limit of synaptic learning, future work should incorporate biologically realistic learning rules ^55-59^.

Together, these findings support a unified view: the cerebellum integrates multiple inhibitory pathways and sparse coding strategies to flexibly adapt its computations to diverse functional demands. Decoding how these mechanisms are balanced will be crucial for advancing models and experiments that deepen our understanding of cerebellar roles well beyond motor control.

## Method Details

### Cerebellar Circuit Model

We constructed a cerebellar circuit model grounded in well-characterized cerebellar anatomy and electrophysiology (Figure 1). The model comprises excitatory mossy fibers, excitatory granule cells, inhibitory Golgi cells, and inhibitory Purkinje neurons. Granule cell activity is regulated by inhibitory inputs from Golgi cells. All synaptic connections are fixed except for the plastic granule cell→Purkinje neuron synapses, which adapt to reproduce or identify target traces and patterns by minimizing error signals. The model code will be made publicly available upon acceptance of the manuscript.

The model structure follows prior work ^29,47^. Each granule cell receives *M*_1_ synaptic inputs from a randomly selected subset of mossy fibers. The activity of the *j*_*th*_ granule cell is given by:

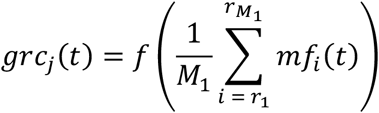

where *f* (*x*) = max (*x*, 0) is a rectified linear activation function.

### Inhibition via FFI and FBI Pathways

Granule cell activity is shaped by Golgi cell inhibition, activated either by mossy fibers (FFI) or granule cells (FBI). Assuming linear computations ^60^:

### FFI Pathway

The mean postsynaptic excitation in Golgi cells from mossy fibers is:

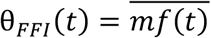

Granule cell activity is then:

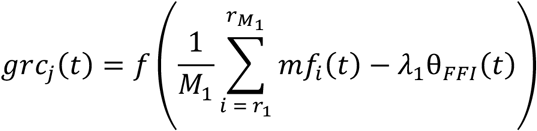

### FBI Pathway

The *k*_*th*_ Golgi cell receives *M*_2_ excitatory inputs from random granule cells:

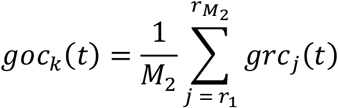

The *j*_*th*_ granule cell receives *M*_3_ inhibitory inputs from random Golgi cells:

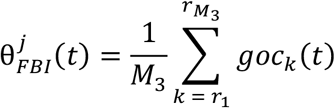

Twenty precents of Golgi cells are randomly inactivated. Granule cell activity is:

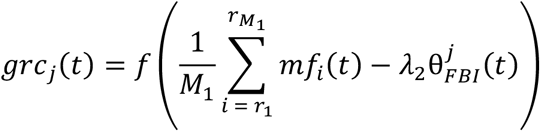

### Combined Inhibition

The net synaptic drive is the mossy fiber excitation minus a weighted sum of FFI and FBI inhibition:

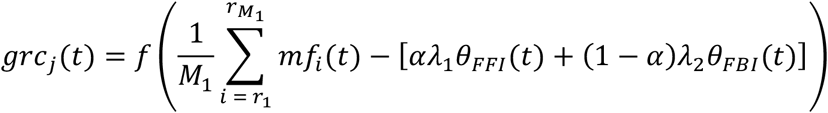

Here, *λ*_1_ and *λ*_2_ are inhibitory gains shared across granule cells, and *α* controls the relative contribution of each pathway.

By assuming linear computations in Purkinje neurons ^60,61^, the activity of the *l*_*th*_ Purkinje neuron is:

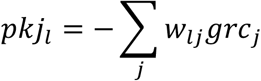

### Synaptic Learning

We implemented a supervised gradient descent rule for granule cell→Purkinje neuron synapses:

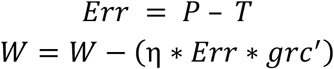

where *P* is the Purkinje neuron output, *T* is the target trace or pattern, *W* is the synaptic weight matrix, and η is the is the learning rate.

### Synaptic Consolidation for Incremental Learning

We incorporated two biologically motivated mechanisms: Elastic weight consolidation (EWC) ^36^ and synaptic intelligence (SI) ^37^.

EWC penalizes changes to weights important for prior tasks, quantified by the Fisher Information Matrix *F*:

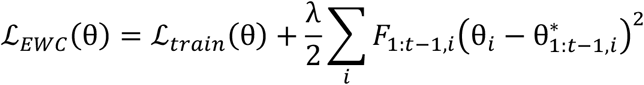

SI assigns local importance values based on past contributions to loss reduction:

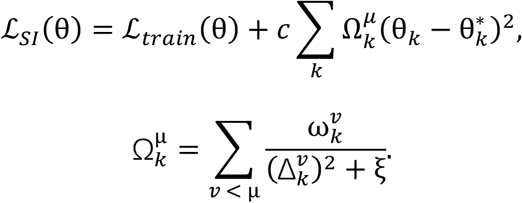

### Activation Level Metric

Granule cell activation level is defined as the spatio-temporal average fraction of active neurons:

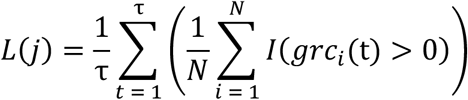

where *N* is the number of granule cells, τ is the stimulation duration, and *I*(·) is the indicator function that outputs 1 when a cell’s activity is non-zero and 0 otherwise.

### Task Protocols

#### Complex Trace Learning

Auditory signals from the TI-46 corpus ^34^ were converted into 12-channel cochleograms as mossy fiber inputs. Each digit (“0”–”9”) was mapped to a three-dimensional trajectory (*x, y, z*). To assess robustness, Gaussian noise 𝒩(0,0.01) was added to 20% of temporal coordinates.

Sequential learning introduced digit-specific mappings progressively, with performance measured by average MSE across tasks:

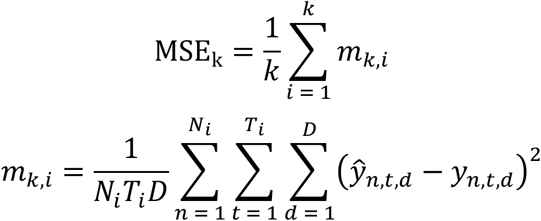

where *N*_*i*_ is the number of test samples in task *i, T*_*i*_ is the sequence length, and *D* is the dimensionality of the output. In this setting, *D* = 3, corresponding to each trajectory’s three coordinates (*x, y, z*).

This metric captures how accurately the network reproduces previously learned trajectories while adapting to newly introduced mappings, providing a reliable measure of continual learning performance in the trace prediction setting.

#### Pattern Identification

MNIST images ^35^ were flattened into 784-dimensional mossy fiber input vectors. Ten Purkinje neurons represented digit patterns (“0”–”9”), with the output determined by the neuron with the highest activity. Sequential learning introduced patterns incrementally, with accuracy measured as:

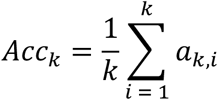

where *a*_*k,i*_ denotes the test accuracy for class *i* after completion of training up to task *k*. This metric captures how well the network retains previously acquired pattern representations while incorporating newly introduced ones, providing a reliable measure of incremental learning performance across sequential training stages. In the interleaved learning condition, all data were presented to the model in an interleaved fashion.

## Acknowledgements

This work is supported by the National Key Research and Development Program of China (2023YFF1204200) and the National Natural Science Foundation of China (12372060, 62476197).

## Supplementary Figures

**Figure S1.**
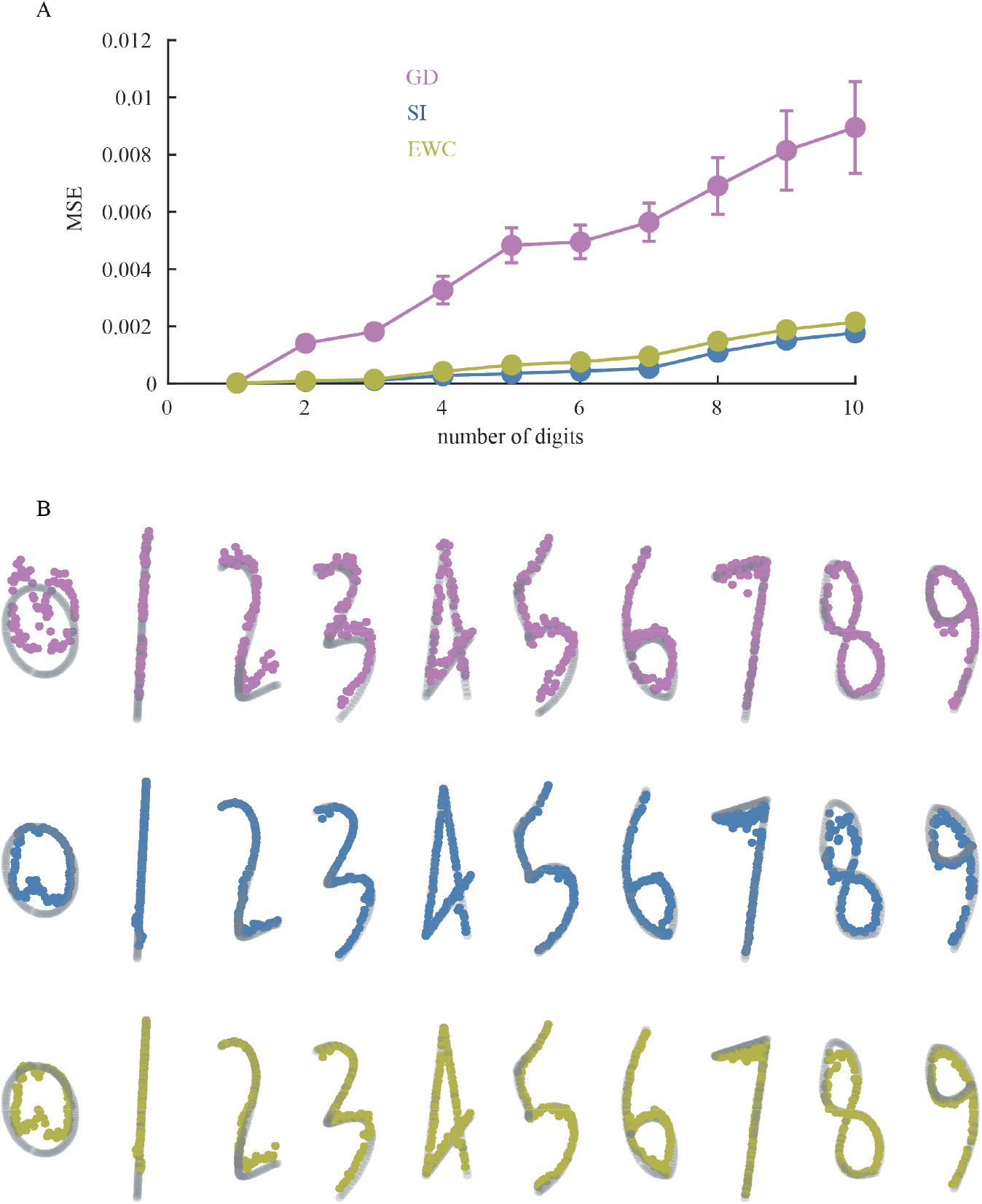
Impact of Advanced Synaptic Update Strategies on Incremental Learning of Complex Digit Traces. (A) Mean squared error (MSE) of incremental digit trace learning as a function of the number of digits learned. Color indicates different synaptic update strategies. Gradient descent is the default synaptic update method. Synaptic intelligence and elastic weight consolidation are advanced strategies. They represent the upper bound of biologically plausible synaptic update mechanisms. (B) Optimal incremental digit trace learnings achieved with different synaptic update strategies. Panels from top to bottom show results for gradient descent, elastic weight consolidation, and synaptic intelligence, respectively.

**Figure S2.**
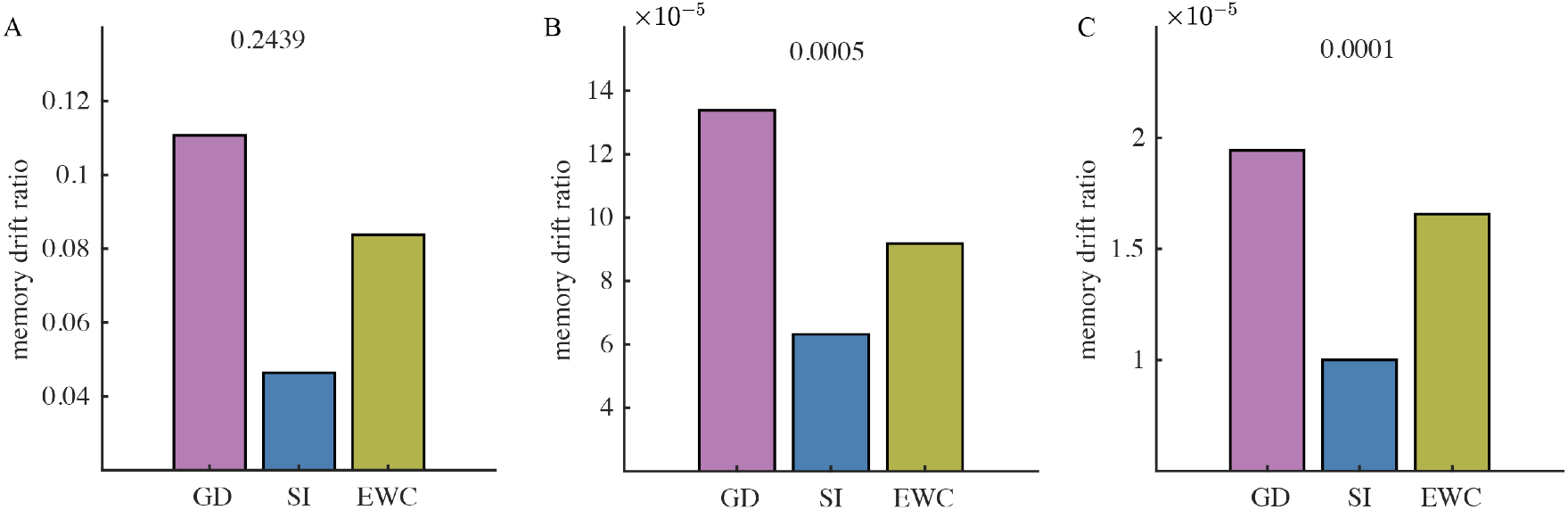
Impact of Advanced Synaptic Update Strategies on Reducing Memory Drift Ratio in Digit Trace Incremental Learning. (A-C) Effects of synaptic intelligence and elastic weight consolidation on reducing the memory drift ratio when granule cells are activated at levels of 0.2439, 0.0005, and 0.0001, respectively. Note: The vertical axis scales differ across panels.

**Figure S3.**
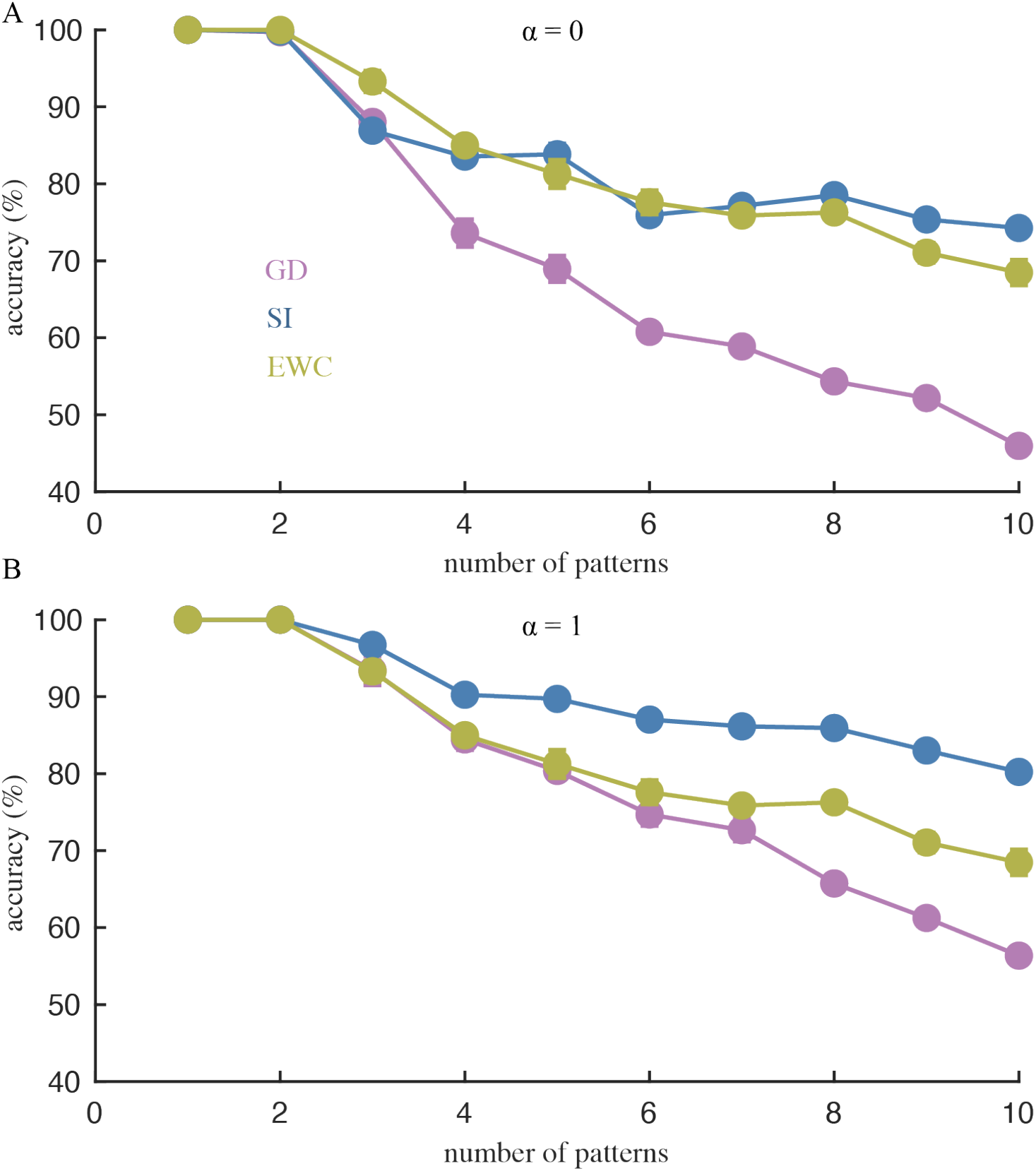
Impacts of Advanced Synaptic Update Strategies on Incremental Pattern Identification. (A) The identification accuracy as a function of the number of patterns incrementally learned in the model regulated by the pathway of FBI. (B) The identification accuracy as a function of the number of patterns incrementally learned in the model regulated by the pathway of FFI.

**Figure S4.**
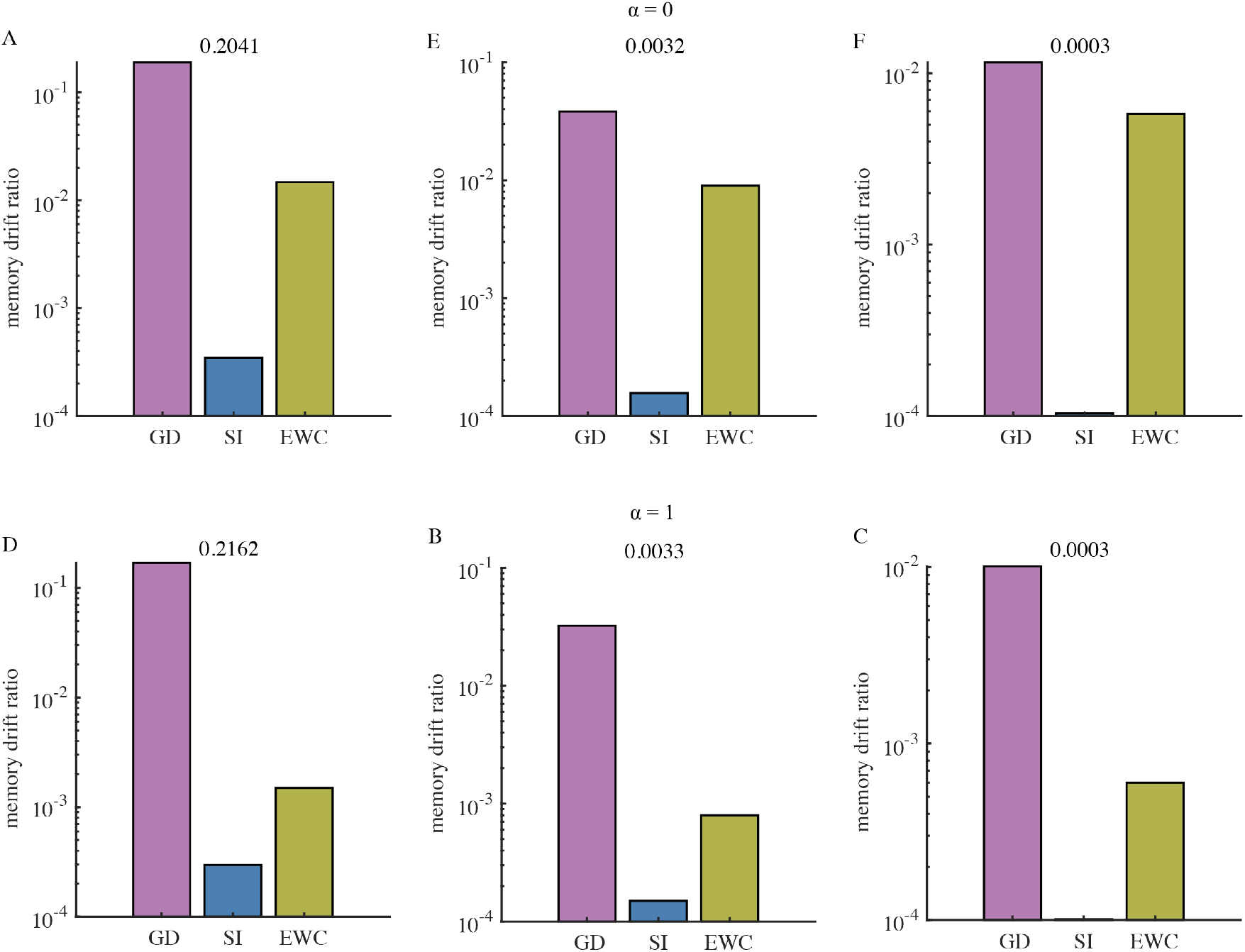
Impacts of Advanced Synaptic Update Strategies on Reducing Memory Drift Ratio in Incremental Pattern Identification. (A–C) Effects of synaptic intelligence and elastic weight consolidation on reducing the memory drift ratio when granule cells are activated at levels of 0.2041, 0.0032, and 0.0003, respectively, in the model regulated by the FBI pathway. (D–F) Same as (A– C), but in the model regulated by the FFI pathway. Activation levels differ slightly, with values of 0.2162, 0.0033, and 0.0003, respectively. Note: Vertical axis scales vary across panels.

